# Tibetan antelope rests like a Puppet

**DOI:** 10.1101/413393

**Authors:** Yunchao Luo, Lin Wang, Le Yang, Ming Tan, Yiqian Wu, Yuhang Li, Zhongqiu Li

## Abstract

Rest contributes to a large part of animals’ daily life, and animals usually rest in two ways, standing or in recumbence. For small or medium sized ungulates, they bed to rest in most cases, and standing rest is very rare and hardly seen. Here we described a standing rest behaviour of medium sized Tibetan antelopes (*Pantholops hodgsonii*) living on the roof of the world, Tibet Plateau, which has not been reported before. We named the standing rest behaviour here as Puppet behaviour, since the antelope can stand still for a certain time just like a Puppet. Of the total 304 focal individuals, 48.3% (98/203) of adult and sub-adult males expressed the Puppet behaviour, whereas only 6.3% (6/96) of females did, indicating an obvious sexual difference. Puppet behaviour occurred more frequently at noon and in the afternoon on sunny and cloudy days, meaning that day time and weather were both influential factors. Puppet behaviour was usually accompanied with rumination and sometimes ended with leg-shaking. Our results suggest that Puppet behaviour is probably an adaptive form of rest, which serves a thermoregulatory and anti-predation function, and is much simpler and safer than recumbent rest.

---

Animals usually behave in a relatively fixed manner, and these common behaviours can be classified into several categories, including feeding, resting, moving, alerting, grooming, etc. Animals need rest or sleep to perform a number of physiological functions such as saving energy, thermoregulation, and maintenance of their immune system [1, 2]. Generally, animals need to sleep or take rest for 2 to more than 20 hours a day to recover from the exhaustion due to daily activities [3-6]. Even during the daytime, animals spend a large amount of their time to rest [3].

Rest is a perilous condition of reduced activity, little consciousness, greatly reduced responsiveness, during which the natural enemies can hunt the resting animals [7, 8]. Therefore, animals need to adopt a proper way and time to rest, in order to recover physiologically and avoid their predators. Animals take rest either by standing or bedding (in recumbence) [9]. Standing rest highlights the animal as an obvious predation target that can easily be observed by the natural enemies [7]. Thus to adopt standing rest probably depends upon the anti-predator ability of the animals or more specifically their body size. For small or medium sized cloven-hoofed animals, they prefer bedding to rest. For example, Przewalski’s gazelle (*Procapra przewalskii*) with a body size of about 26 kg, spend 38% of their day time to rest in recumbence [10]. When they stand, they are usually dealing with feeding or alert, and we had never seen their rest while standing. This is also true in other small or medium sized ungulates including Tibetan gazelle (*P. picticaudata*) [11], Asiatic ibex (*Capra sibirica*) [12, 13], and goitred gazelle [14]. However, standing rest seems common in large body-sized ungulates, just like Asiatic wild ass (*Equus hemionus*), they spend nearly one third to rest [14]. Similarly, elephants have an average daily total sleep (standing or in recumbence) time of 2 h due to the large body size, and only exhibit recumbent sleep every third or fourth day [15-17].

We started our field behavioural study on medium sized Tibetan antelopes in 2016, and found that Tibetan antelopes especially males sometimes kept standing for several min (see supplemental video 1 and Fig 1). During this time, they didn’t feed, didn’t move, and just kept their body motionless, like a puppet. In small or medium sized ungulates, standing usually severs as anti-predation vigilance [11, 18], but this standing-still behaviour or called Puppet behaviour seemed very different from vigilance, since their head also kept motionless and was usually under shoulder, and thus probably cannot detect a whole of their surroundings.

**Figure 1.**
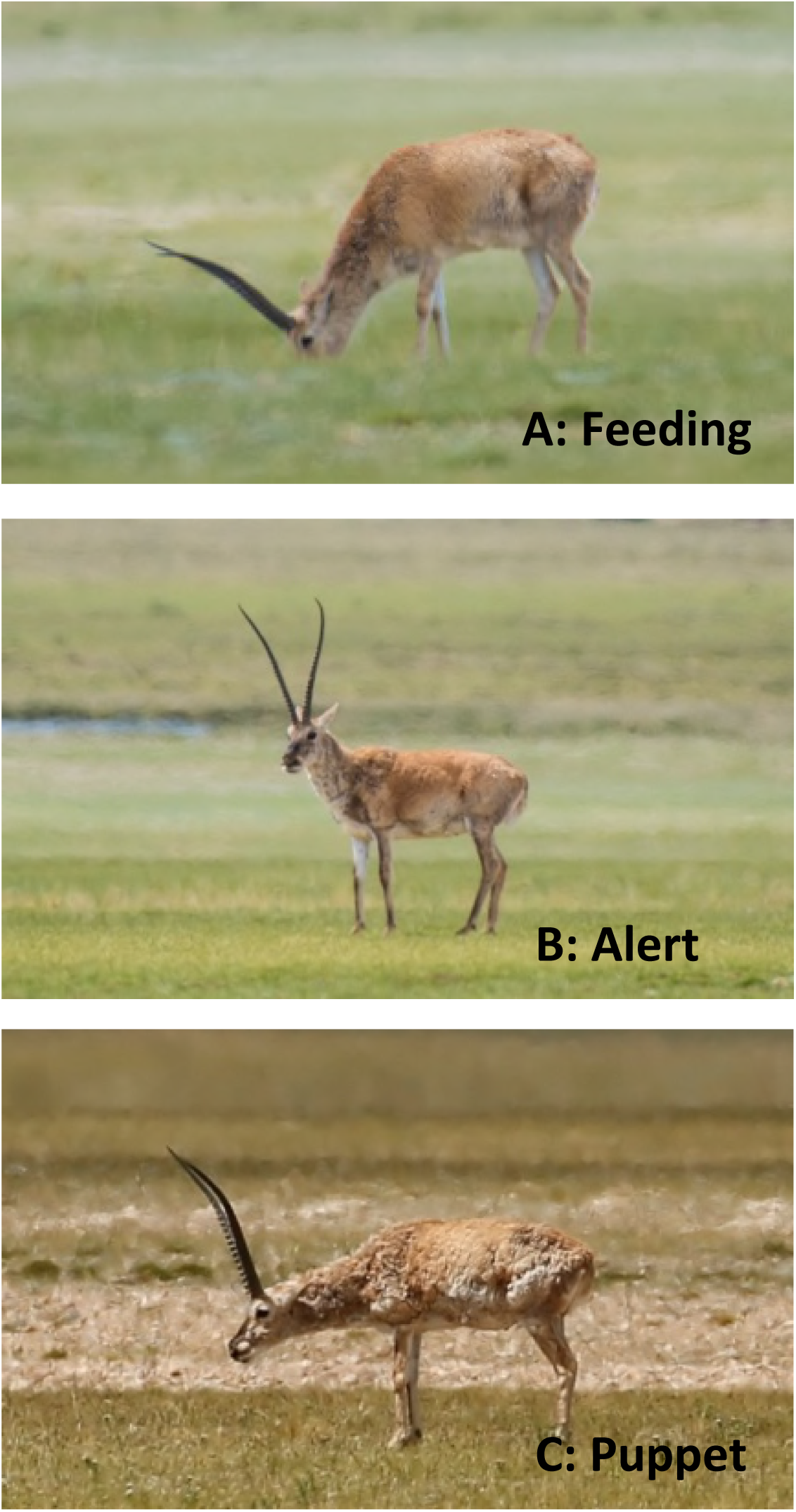
Photos of feeding (A), vigilance (B) and Puppet behaviour (C) of Tibetan antelopes

To our knowledge, there is not any report on this kind of standing behaviour in Tibetan antelopes, or other median or small sized ungulates. We considered that the Puppet behaviour serves a primary function of rest, thus we can predict that this behaviour would occur less frequently in early morning, since they need more time to feed after a whole night rest. Puppet behaviour might also serve a thermoregulatory function, because when they stand still, they have a much larger body surface contact with the air, and therefore we predict that the Puppet behaviour would occur more frequently on sunny or cloudy days than on rainy or overcast days. Puppet behaviour means motionless, and can be considered as a vulnerable time for the antelopes but a great time for predators. As a sexually segregated species, males of Tibetan antelope are much larger and stronger, and thus more tolerant than females when confronting with predators. From a view of anti-predation, we can predict that Puppet behaviour would occur more frequently in males than in females. Here we collected two years data to explore if the occurrence of Puppet behaviour is related to the day time, weather and sex-age.

## Materials and Methods

### Study areas

This study was conducted in Shenzha County (30°02′39 ″~32°19′33 ″ N, 87°45′30″~89°47′49 ″ E), which is located in the central part of Tibet Autonomous Region, China. Elevations range from 4,530m to 6,448m, with an average of 4700m. Local climate is characterized by extreme cold and long winters, strong winds and high levels of solar radiation. Mean annual temperature was 0.4°C. Annual precipitation is about 330 mm and most rain falls between June and September. Alpine meadow is the main vegetation type and no shrubs appear in this area.

### Study species

The Tibetan antelope is a flagship species to Qinghai-Tibet Plateau. The total population underwent a severe decline in the 1980s and early 1990s as a result of commercial poaching for the valuable underfur [19]. Rigorous protection has been enforced since then, and recent estimates were made at about 100,000 ~ 150,000 [20, 21]. Tibetan antelope has been classified as Endangered by IUCN since 2000 and is a Category I (Endangered in China) National Protected Wild Animal Species in China since 1989. Tibetan antelope breeds from December to January and the lambing season is from June to July. They are dimorphic; adult male with an average body weight of 39kg, is larger than female with an average of 26 kg [21]. The resident status of Tibetan antelope can be divided into two types, the migratory population and the resident population. Our focal population in Shenzha is resident and does not migrate. The population might be up to 10, 000.

The most significant mammalian predator in this areas is the wolf (*Canis lupus*), which is relatively common; the snow leopard (*Panthera uncia*), the lynx (*Felis lynx*) and the brown bear (*Ursus arctos*), which are much rarer. Tibetan fox (*Vulpes ferrilata*), are also common and may prey upon lambs of the antelopes. Large raptors including upland buzzard (*Buteo hemillasius*), cinereous vulture (*Aegypius monachus*) and lammergeier (*Gypaetus barbatus*) are common and are frequent scavengers of dead ungulates.

### Behavioural sampling

Daytime observations were carried out from sunrise to sunset in the summer (July & August of 2016, June & July of 2017) in Shenzha. We defined a group as a herd of antelopes with no more than 50 m separating any two group members. Observations were carried out from the roadside using binoculars (8X42) or a telescope (20~60X63). We walked or drove regular routes to find targets for video recording.

For most groups, we focal sampled one or two individuals. We may collect a few more samples for some large female groups (usually more than a few hundred individuals), but we will collect from different part of the group to avoid resampling. It was not practically feasible to mark individuals or to recognize individuals through particular features, and thus there was a possibility that a focal individual was recorded more than once. However, since the population was more than 10 000 and extremely large, the possibility of resampling in a same day was rather small.

At the beginning of each focal observation, we recorded the date, time of day, location, weather, group type, and group size. Since the antelopes are sexually segregated, only three group types (single-male groups, single-female groups and mother-lamb groups) could be found during summer. Focal individuals were classified into four categories: adult male, sub-adult male, female and lamb. It was not practically feasible to distinguish adult females from sub-adult females, and thus the two age classes were therefore collapsed.

Behavioural events were video recorded. Observations lasted 30 min unless we lost sight of the focal individual. Puppet behaviour is defined as standing still, head up or down, and without any other apparent acts.

### Statistical Analysis

We collected a total of 304 behavioural samples, and the total observation time was 3968 min with an average (± SE) observation length of 13.1 ± 0.2 min. Since there were not any Puppet behaviours expressed in more than half of these samples, we firstly established a logistic regression model, to see if any effects of day time (morning: before Beijing Time 13:00 (local time 11:00); noon: Beijing time 13:00-16:00 (local time 11:00-14:00); afternoon: after Beijing time 16:00 (local time 14:00)), weather (sunny, cloudy, overcast or rainy), sex-age (adult male, sub-adult male, female, and lamb), and group size (1-400 with a median of 7) on the occurrence of Puppet behaviour. Then we kept samples expressed at least one time of Puppet behaviour (104 samples left), and established a general linear model, to see if the duration of Puppet behaviours respond to day time, weather, sex-age, and group size. All these analysis were done using SPSS 19.0, values were shown with Mean ± SE, and significant levels were set at 0.05.

## Results

The Puppet behaviour was expressed 344 times in 104 of the total 304 samples. Hereafter we described the behaviour with respect to each environmental and social factor.

Puppet behaviour was defined as standing still, without any acts of their whole body for a certain time (ranged from 2 to 842 s, with a median of 57 s). We measured the angle between their neck and foreleg, and found that the angle was usually between 40° and 100° (Fig 1 & 2). Comparably, when the antelopes were feeding, the angle was usually less than 40°; and when they were in vigilance, the angle was always more than 120°, with head up and scanning around meanwhile. 63.9% (92/144) of the clearly observed Puppet behaviours were found to be accompanied by oral rumination (supplemental video 2). And when the antelopes ended the Puppet behaviour, sometimes (102/344, 29.7%) they would shake their hind legs.

**Figure 2.**
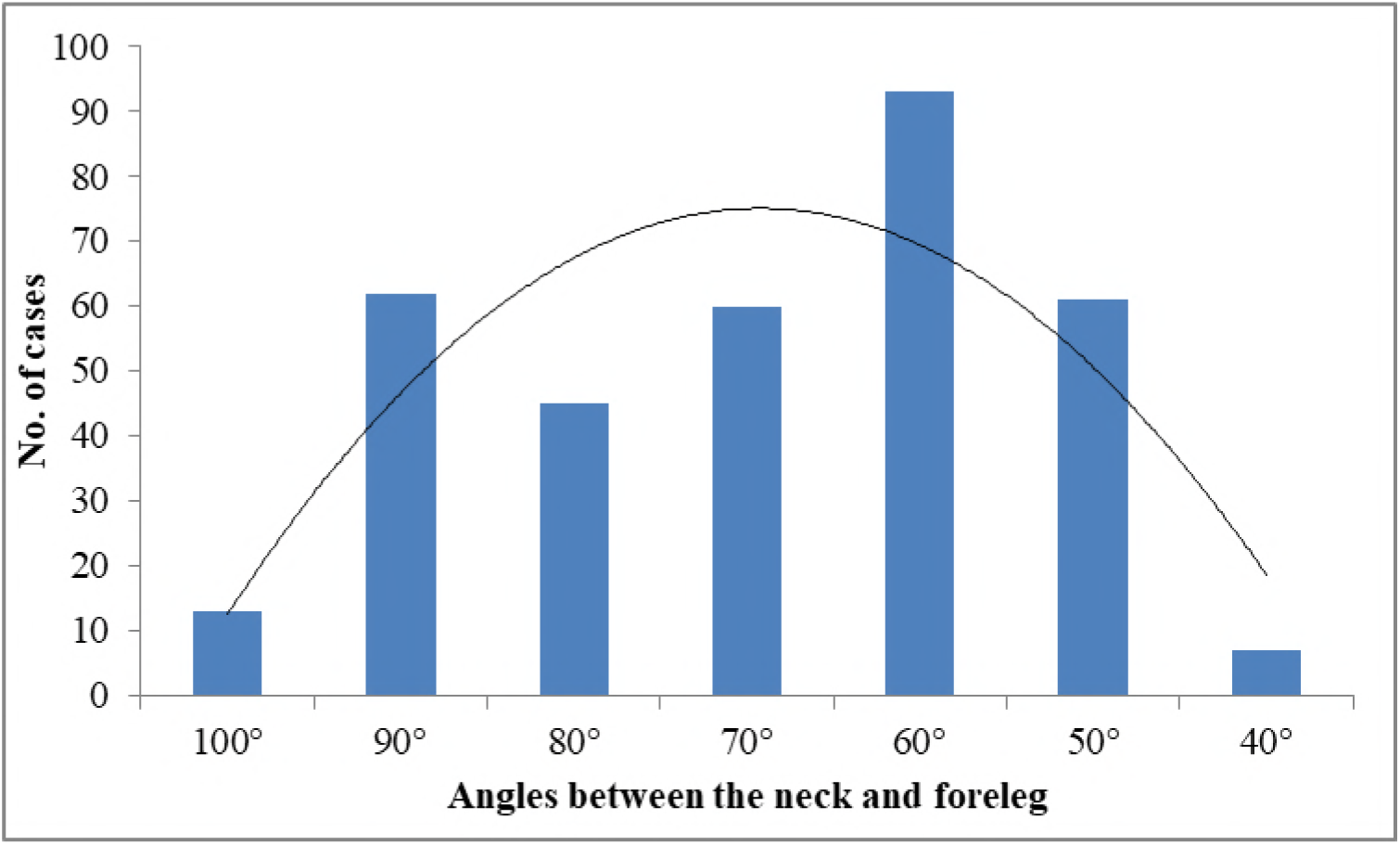
Posture expression of Puppet behaviour of Tibetan antelopes

With the help of the logistic model, we found that the occurrence of Puppet behaviour was affected by day time (Wald=8.114, df=2, P=0.017, fig 3), weather (Wald=22.121, df=2, P<0.001, fig 4), and sex-age (Wald=23.984, df=3, P<0.001, fig 5), but not group size (Wald=1.279, df=1, P=0.258). Compared to the afternoon (28/61, 45.9%), Puppet behaviour occurred less in the morning (27/137, 19.7%; B=-1.261±0.461, Wald=7.500, df=1, P=0.006) but not noon (49/106, 46.2%; B=-0.621±0.455, Wald=1.864, df=1, P=0.172). Compared to overcast or rainy days (6/64, 9.4%), the antelopes expressed much more Puppet behaviours on sunny days (30/52, 57.7%; B=2.404±0.545, Wald=19.486, df=1, P<0.001) or cloudy days (68/188, 36.2%; B=2.119±0.487, Wald=18.949, df=1, P<0.001). Compared to adult females (6/96, 6.25%), adult males expressed much more Puppet behaviours (93/177, 52.5%; B=2.503±0.572, Wald=19.133, df=1, P<0.001), sub-adult males expressed a little more but not significantly (5/26, 19.2%; B=1.103±0.759, Wald=2.115, df=1, P=0.146), and lambs did not express any Puppet behaviour (0/5, 0%).

**Figure 3.**
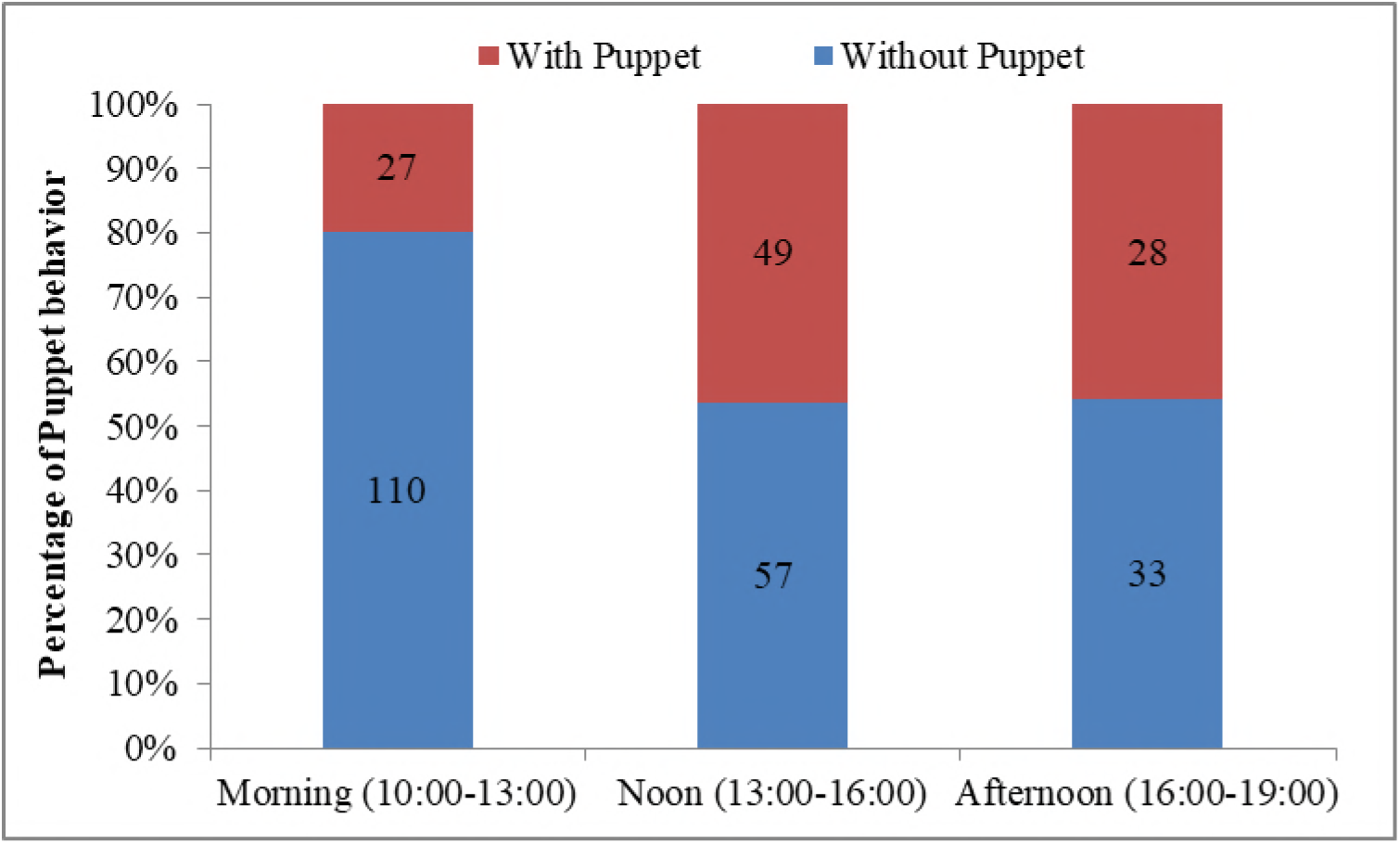
Effect of day time on the occurrence of Puppet behaviour of Tibetan antelopes

**Figure 4.**
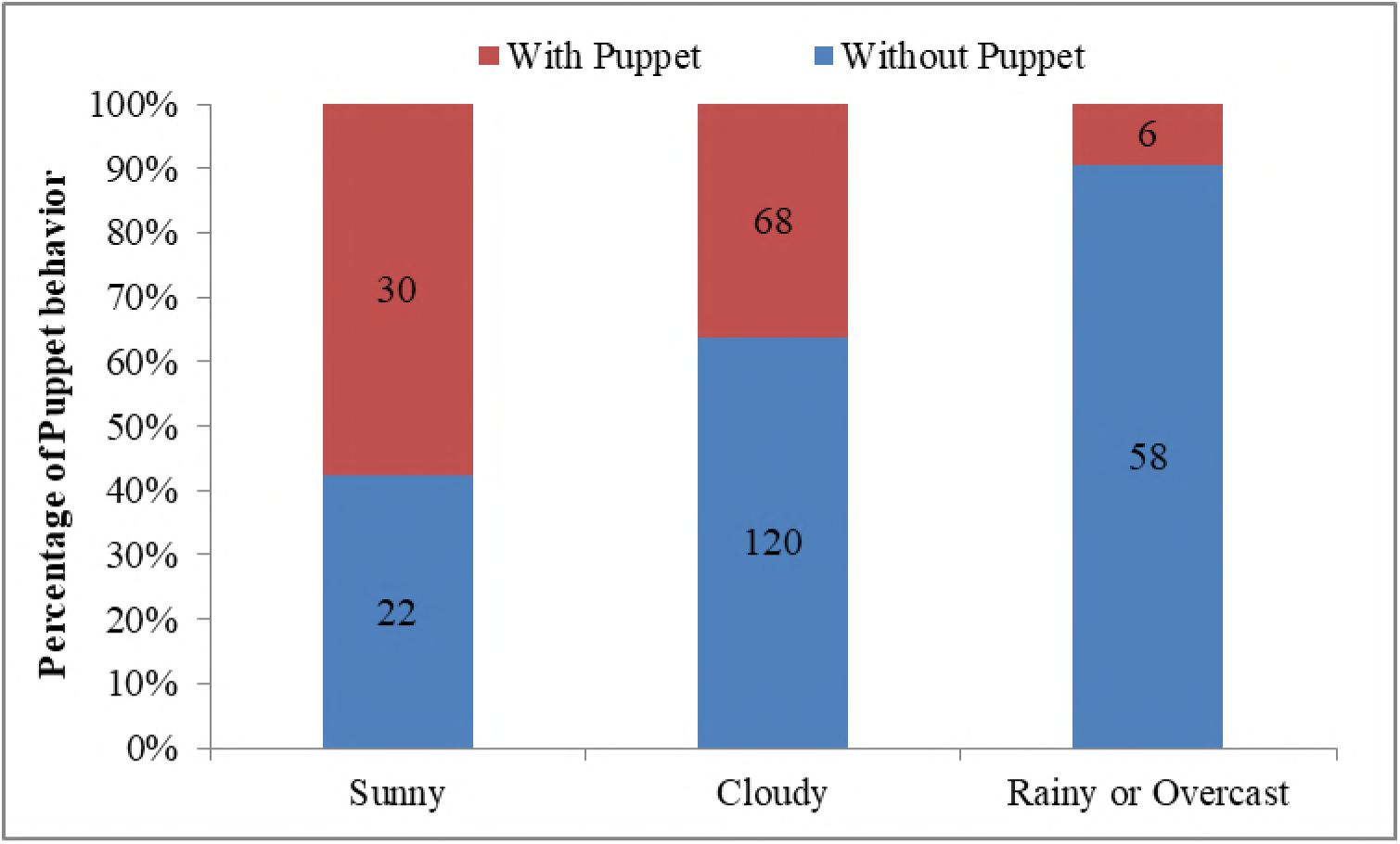
Effect of weather on the occurrence of Puppet behaviour of Tibetan antelopes

**Figure 5.**
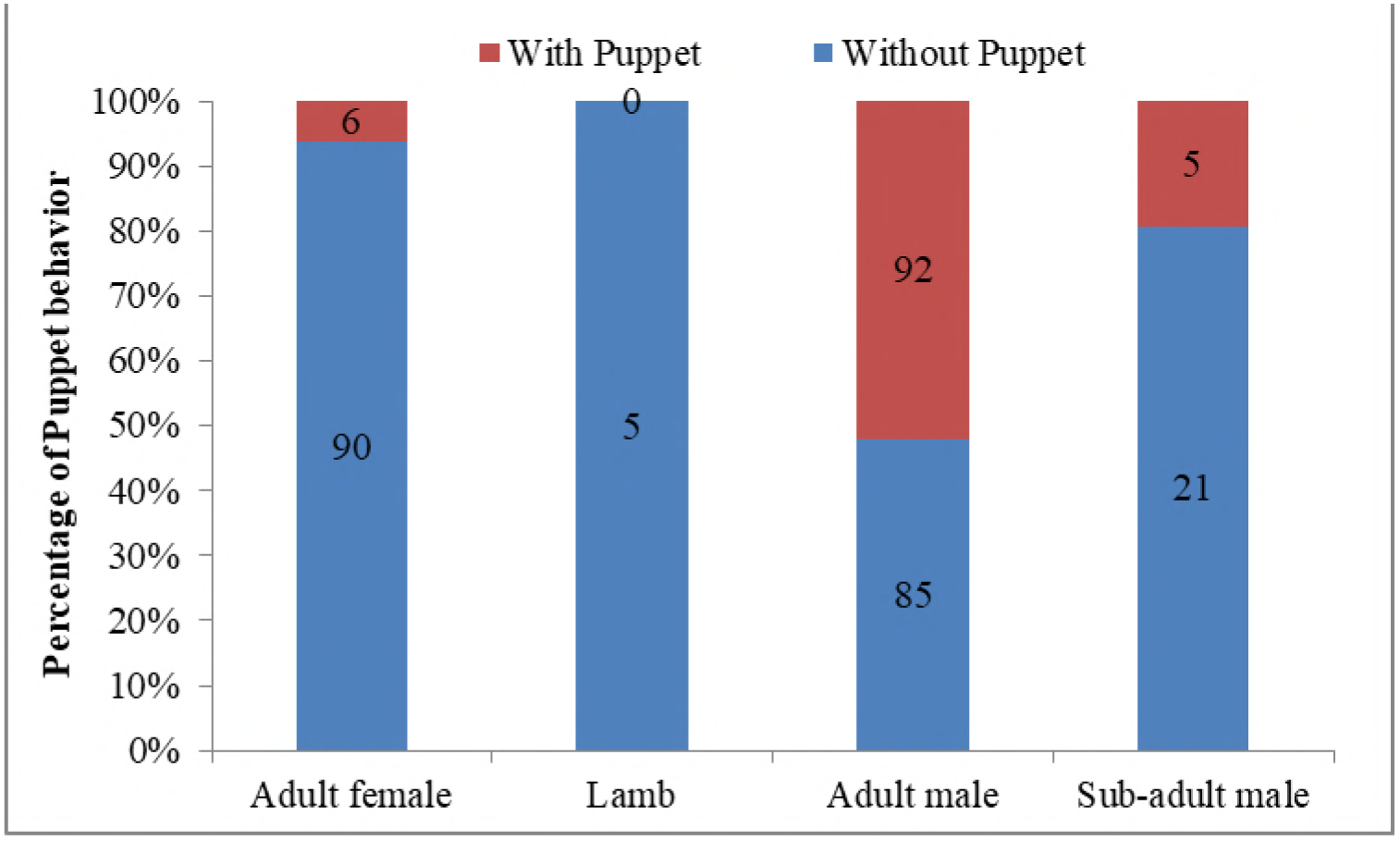
Effect of sex-age on the occurrence of Puppet behaviour of Tibetan antelopes

Similarly, we established another logistic model and found that the occurrence of recumbent rest was independent of time (Wald=1.726, df=2, P=0.422), sex-age (Wald=0.250, df=2, P=0.969), group size (Wald=1.974, df=1, P=0.160), but marginally influenced by weather (Wald=5.962, df=2, P=0.051). Compared to the rainy day, the antelopes bedded more on sunny days (B=0.911±0.447, Wald=4.159, df=1, P=0.041).

With the help of general linear model, we explored if any factors had an effect on the duration of Puppet behaviour. The duration of Puppet behaviour in females lasted 42.7±41.5 s, much shorter than that of adult males (91.5±14.1 s) or sub-adult males (100.3±43.3 s), however, the difference was insignificant (F_2,96_=0.736, P=0.482). Similarly, we did not find any significant effect of weather (F_2,96_=0.429, P=0.652), day time (F_2,96_ =2.462, P=0.091) and group size (F_1,96_ =0.035, P=0.851) either.

## Discussion

Our result indicates that the Puppet behaviour of Tibetan antelope occurred more frequently at noon and in the afternoon, on sunny and cloudy days, and in males, indicating that all the three factors (day time, weather, and sex-age) shaped the expression of Puppet behaviour. Hereafter we discussed the functional explanations for the Puppet behaviour.

Animals rest and sleep in their own way, bedding or standing, and whether to select standing rest is probably depending on their body size. Generally large body-sized ungulates rest in both standing and recumbent ways, and the standing rest can contribute to an even larger part than recumbent rest, e.g, Przewalski’s horses (*E. przewalskii*) spent 15.7% of a whole day and up to 25% of their diurnal time in standing rest [22]; Giraffes spent about 20% of their time in recumbent rest, but standing was also a very important way of rest, in mothers, standing rest even contributed to 1/3 of their night time from 4:00 to 7:00 [9]. However, standing rest seems very rare in median or small sized ungulates, and recumbence is probably the main or the only way to rest, as we found in other Tibetan ungulates [10]. As a median ungulate, the frequent expression of standing rest by Tibetan antelopes, especially by males, seems difficult to interpret.

We proposed two possible explanations, energy saving and anti-predation, both closely related to body size. With the increase of body size, the rest progress from standing to recumbence and then back to standing, this could be costly in terms of both time and energy. This extremely slow progress is very risky and may lead to a fatality in the case of an immediate attack. Standing rest for a large body-sized animals indicate them as obvious targets for the predators to find them easily, even from a long distance. However, they have larger body sizes and are more tolerant ofpredators; the risk of being preyed on may be lower for them as compared to smaller animals. And due to standing with a better visual field, they can probably find the predators earlier and react timely. In Tibetan antelopes, body size principle also works. As expected, larger males of the Tibetan antelope exhibited standing rest for much more times than smaller females’ counterpart. Body size, as well as its associated ability of anti-predation, probably becomes the determinant why males preferred standing rest.

Another point is, animals with larger body size would probably have evolved an adaptive body structure to keep balance and save energy while standing rest [23]. Just like horses, the skeleton is helpful to support and balance themselves when they are resting with standing state [23]. For Tibetan antelopes, we consider that the standing state might be a relaxed way to rest because the angle between their neck and foreleg was around 70°, not like vigilance behaviour with an angle of more than 90°or feeding behaviour with an angle of less than 40°, both of which need an intense coordination of both skeletal and muscular systems. This posture of Tibetan antelope is in congruence with the standing rest of flamingos by single leg[24], which needs more mechanical analysis.

Both day time and weather can influence Puppet behaviour, but they were not the determinants since the behaviour also occurred in the early morning and on rainy or overcast days. It is easy to understand that the antelopes expressed a lower Puppet behaviour in the early morning since they have spent a whole night sleep and need more time to feed. The rest starts after they fulfill their stomachs at noon and occurs frequently in the afternoon. The Puppet behaviour occurs more frequently when the weather is sunny or cloudy, and probably it serves a function of thermoregulation. The elevation in this area is very high and the average temperature is relative low compared with other areas [25]. However, the sun radiation is rather strong and the real-time temperature reached 20°C during sunny days. While standing, the surface area of the bodies of the antelopes is larger, they can have much body contact with the air, which can facilitate heat dissipation. This is also true in other ungulates, such as Asiatic wild ass [26], Przewalski’s horse [27, 28], African wild ass (*E. africanus*) [29] and Grevy’s zebra (*E. grevyi*) [30].

The Puppet behaviour of Tibetan antelope is like a kind of REM (rapid eye movement) sleep which is a state with many neural and physiological activities [31, 32]. The duration of the standing rest ranged from one or two min, and most of the times (63.9%; 92/144) standing rest was accompanied by oral rumination. Rumination can be considered as a continuation of feeding behaviour, and it usually occurs while rest, no matter standing or in recumbence [33]. Since ungulates need a large amount of vegetarian food a day, e.g., an adult giraffe consumes about 37 kg of food per day [9], thus they would spend a large amount of daily time in ruminating. Actually EEG recordings have shown that rumination is not exclusive with rest [9]. REM sleep or rest at a standing posture may be relaxing for the antelope to ruminate their food and recover their body, as well as to be ready to escape should an attack occur [7, 32]. However, this assumption opens a new window to future research for confirmation.

Leg-shaking behaviour was also found sometimes when the antelopes wanted to end the Puppet behaviour. We considered it as relaxing since a long time keeping body motionless would probably lead to leg numbness. This leg-shaking behaviour is similar to the relaxed rolling behaviour of horses, which also occurs after sleeping [34, 35]. Another research of beluga whale (*Delphinapterus leucas*) also reported that muscle jerks occurred more frequently at the end of rest episodes [36].

## Conclusion

In conclusion, we reported a strange but interesting Puppet behaviour in Tibetan antelopes. This kind of standing rest behaviour can be found more frequently in large sized ungulates but occurred in medium sized Tibetan antelopes. Several factors including sex-age, day time and weather can influence the expression of Puppet behaviour. Further studies should focus on the mechanical analysis on the standing architecture to explore how the antelopes balance themselves while rest.

## Acknowledgment

We sincerely thank the National Natural Science Foundation of the People’s Republic of China (No. 31360141, No. 31772470, and No. J1103512) and West Light Foundation of Chinese Academy of Sciences (2015) for supporting this study financially. We thank Dr. Xinming Lian (Northwestern Plateau Institute of Biology, Chinese Academy of Sciences) for his valuable suggestions, and Gesang, Jinxuan Shen, and Jiabu for their help in the field work and Sana Ullah for language polishment.

## Author Information

### Authors Contributions

YL, LY & ZL designed the study; YL, LY, MT, YW & YL collected the data; YL, LW & ZL carried out data analysis, drafted the manuscript.

### Ethical Approval and Consent to Participate

This is an observational experiment and all observations were made at a distance of more than 200 m. All the experiment procedures in this study were approved by the Chinese Wildlife Management Authority.

## References

1. Schmidt MH. The energy allocation function of sleep: a unifying theory of sleep, torpor, and continuous wakefulness. Neurosci Biobehav Rev. 2014;47:122–53.

2. Opp MR. Sleeping to fuel the immune system: mammalian sleep and resistance to parasites. BMC Evol Biol. 2009;9(1):8–10.

3. Elgar MA, Pagel MD, Harvey PH. Sleep in mammals. Anim Behav. 1988;36(5):1407–19.

4. Siegel JM. Sleep viewed as a state of adaptive inactivity. Nat Rev Neurosci 2009;10(10):747–53.

5. Lutermann H, Verburgt L, Rendigs A. Resting and nesting in a small mammal: sleeping sites as a limiting resource for female grey mouse lemurs. Anim Behav. 2010;79(6):1211–9.

6. Campbell SF, Tobler I. Animal sleep: a review of sleep duration across phylogeny. Neurosci Biobehav Rev. 1984; Fall;8(3):269–300.

7. Lima SL. Sleeping under the risk of predation. Anim Behav. 2005;70(4):723–36.

8. Siegel JM. Do all animals sleep? Trends Neurosci. 2008;31(4):208–13.

9. Tobler I, Schwierin B. Behavioural sleep in the giraffe (*Giraffa camelopardalis*) in a zoological garden. J Sleep Res. 1996;5(1):21–32.

10. Li CW, Jiang ZG, Feng ZJ, Yang XB, Ji Y, Chen LW. Effects of highway traffic on diurnal activity of the critically endangered Przewalski’s gazelle. Wildlife Res. 2009;36(5):379–85.

11. Li Z, Jiang Z. Group size effect on vigilance: Evidence from Tibetan gazelle in Upper Buha River, Qinghai-Tibet Plateau. Behav Process. 2008;78(1):25–8.

12. Xu F, Ma M, Wu YQ, Yang WK. Winter daytime activity budgets of Asiatic ibex Capra sibirica in Tomur National Nature Reserve of Xinjiang, China. Pak J Zool. 2012;44(2):389–92.

13. Xu F, Ma M, Yang W, Blank D, Wu Y. Test of the activity budget hypothesis on Asiatic ibex in Tian Shan Mountains of Xinjiang, China. Eur J Wildlife Res. 2012;58(1):71–5.

14. Xia C, Yang W, Xu W, Xu F, Blank D. The characteristics of time budgets of male goitred gazelle in different rutting period stages. Acta Theriol Sinica. 2013;33(2):144–9.

15. Gravett N, Bhagwandin A, Sutcliffe R, Landen K, Chase MJ, Lyamin OI, et al. Inactivity/sleep in two wild free-roaming African elephant matriarchs – Does large body size make elephants the shortest mammalian sleepers? Appl Categor Struct. 2017;12(3):e0171903.

16. Benedict FG, Lee RC. Further Observations on the Physiology of the Elephant. J Mammal. 1938;19(2):175–94.

17. Tobler I. Behavioral sleep in the Asian elephant in captivity. Sleep. 1992;15(1):1–12.

18. Li Z, Jiang Z, Beauchamp G. Vigilance in Przewalski’s gazelle: effects of sex, predation risk and group size. J Zool. 2009;277(4):302–8.

19. Schaller GB. Wildlife of the Tebetan Steppe. Chicago: The University of Chicago Press; 1998.

20. Bleisch WV, Buzzard PJ, Zhang H, Liu Z, Li W, Howman W. Surveys at a Tibetan antelope Pantholops hodgsonii calving ground adjacent to the Arjinshan Nature Reserve, Xinjiang, China: decline and recovery of a population. Oryx. 2009;43(2):191–6.

21. Leslie DMJ, Schaller GB. *Pantholops Hodgsonii* (Artiodactyla: Bovidae). Mamm Spe. 2008;817(817):1–13.

22. Boyd LE, Carbonaro DA, Houpt KA. The 24-hour time budget of Przewalski horses. Appl Anim Behav Sci. 1988;21(1):5–17.

23. Schuurman SO, Kersten W, Weijs WA. The equine hind limb is actively stabilized during standing. J Anat. 2003;202(4):355.

24. Chang YH, Ting LH. Mechanical evidence that flamingos can support their body on one leg with little active muscular force. Biol Lett. 2017;13(5):20160948. doi:10.1098/rsbl.2016.0948. PubMed PMID:28539457; PubMed Central PMCID:PMCPMC5454233.

25. Li L, Fan J, Zhang X, Li X. Relationships between NDVI and Climate Changes in Tibet,China. Mt Res. 2017;35(1):9–15.

26. Liu W, Yang W, Huang Y, Xu W, Lin J, Xia C, et al. Diurnal time budgets and activity rhythm of the Khulan. Arid Land Geogr. 2012;35(4):607–14.

27. Dierendonck MCV, Bandi N, Batdorj D, Dügerlham S, Munkhtsog B. Behavioural observations of reintroduced Takhi or Przewalski horses (*Equus ferus przewalskii*) in Mongolia. Appl Anim Behav Sci. 1996;50(2):95–114.

28. Chen J, Hu D, Li K, Cao J, Meng Y, Cui Y. The diurnal feeding behavior comparison between the realeased and captive adult female Przewalski’s horse (*Equus przewalskii*) in summer. Acta Ecol Sinica. 2008;28(3):1104–8.

29. Moehlman PD. Behavioral patterns and communication in feral asses (*Equus africanus*). Appl Anim Behav Sci. 1998;60(2–3):125–69.

30. Klingel H. Soziale Organisation und Verhalten des Grevy-Zebras (*Equus grevyi*). Ethology. 1974;36(1-5):37–70.

31. Snyder F. Toward an evolutionary theory of dreaming. Am J Psychiatry. 1966;123(2):121–42.

32. Voss U. Functions of sleep architecture and the concept of protective fields. Rev Neurosci. 2004;15(1):33–46.

33. Li Z. Sex-Age Related Rumination Behavior of Père David’s Deer under Constraints of Feeding Habitat and Rainfall. PLoS One. 2013;8(6):e66261.

34. Pedersen GR, Søndergaard E, Ladewig J. The influence of bedding on the time horses spend recumbent. J Equine Vet Sci. 2004;24(4):153–8. doi:10.1016/j.jevs.2004.03.013.

35. Hansen MN, Estvan J, Ladewig J. A note on resting behaviour in horses kept on pasture: Rolling prior to getting up. Appl Anim Behav Sci. 2007;105(1-3):265–9. doi:10.1016/j.applanim.2006.04.032.

36. Lyamin OI, Shpak OV, Nazarenko EA, Mukhametov LM. Muscle jerks during behavioral sleep in a beluga whale (*Delphinapterus leucas L.*). Physiol Behav. 2002;76(2):265–70.

